# Detecting ice artefacts in processed macromolecular diffraction data with machine learning

**DOI:** 10.1101/2021.10.28.466246

**Authors:** Kristopher Nolte, Yunyun Gao, Sabrina Stäb, Philip Kollmansberger, Andrea Thorn

## Abstract

Contamination with diffraction from ice crystals can negatively affect, or even impede macromolecular structure determination and therefore, detecting the resulting artefacts in diffraction data is crucial. However, once the data have been processed, it can be very difficult to automatically recognize this problem. To address this, a set of convolutional neural networks named Helcaraxe has been developed which can detect ice diffraction artefacts in processed diffraction data from macromolecular crystals. The networks outperform previous algorithms and will be available as part of the AUSPEX webserver and CCP4-distributed software.

**Synopsis:** A program utilizing artificial learning and convolutional neural networks, named Helcaraxe, has been developed which can detect ice crystal artefacts in processed macromolecular diffraction data with unprecedented accuracy.

## 1. Introduction

Crystals of biological macromolecules are routinely cryo-cooled to a temperature of 100K before exposure to X-rays to reduce radiation damage during the diffraction experiment (Garman & Weik, 2019). Cryo-cooling can lead to the formation of ice (Garman & Owen, 2006). While anti-freeze agents and flash-cooling are commonly employed to minimise this, rings from the diffraction of small ice crystals are frequently found in diffraction images from cryo-cooled macromolecular samples (Chapman & Somasundaram, 2010) (Fig. 1). The emergence of fast-readout pixel detectors which ideally are used to measure finely sliced diffraction images makes visual identification of ice artefacts from individual images more difficult (Thorn *et al*., 2017). Ice rings can be observed when several images are averaged to increase contrast or through newly developed machine learning approaches (Czyzewski *et al*., 2021).

**Figure 1.**
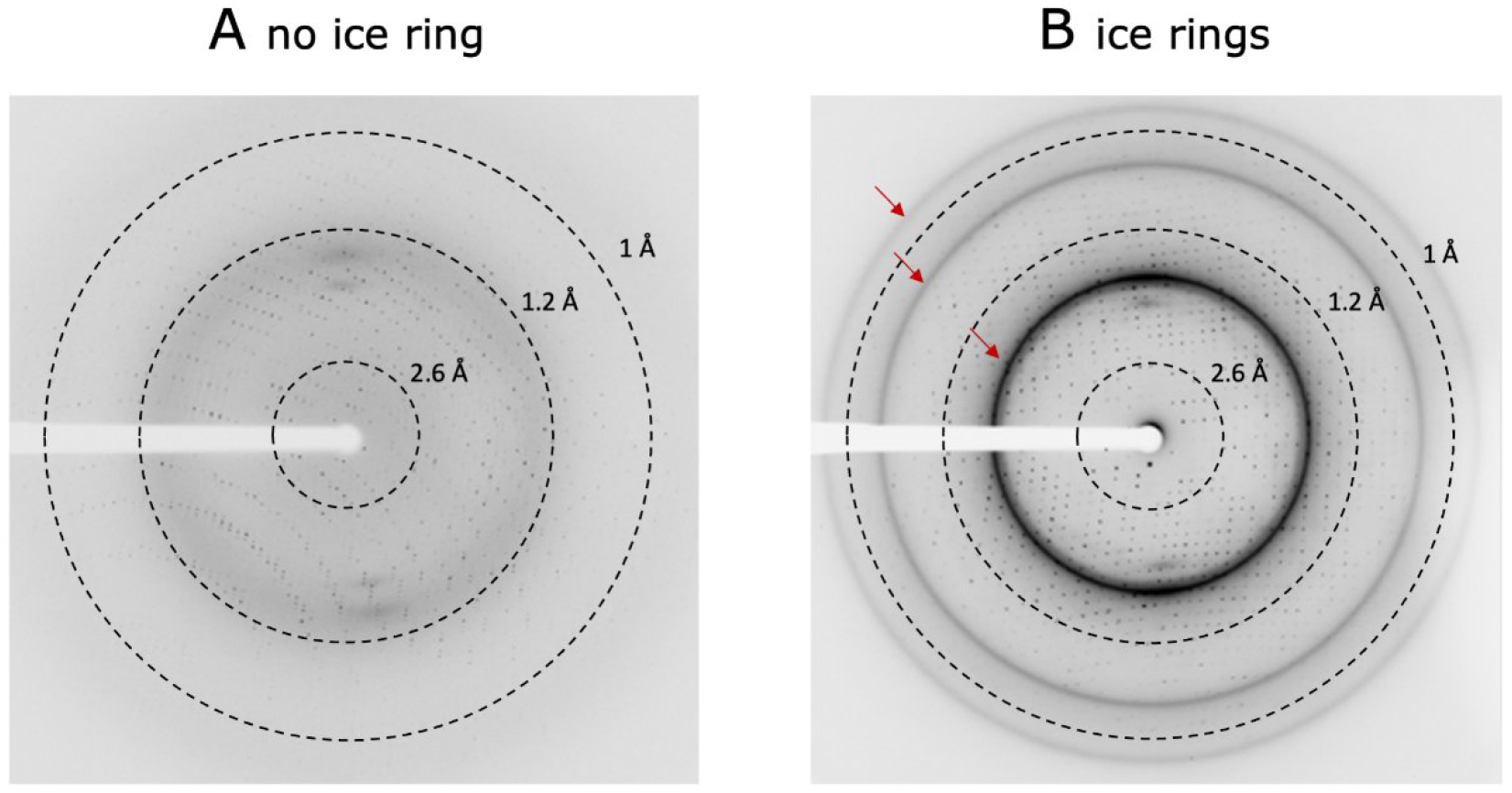
Diffraction patterns of dihydrofolate reductase from mycobacterium ulcerans. The resolution is marked through dashed lines. A shows a pattern without ice rings (PDB-ID: 7K6C) and B with ice crystal artefacts (indicated by the red arrows) (PDB-ID: 7KM9).

**Figure 1.**
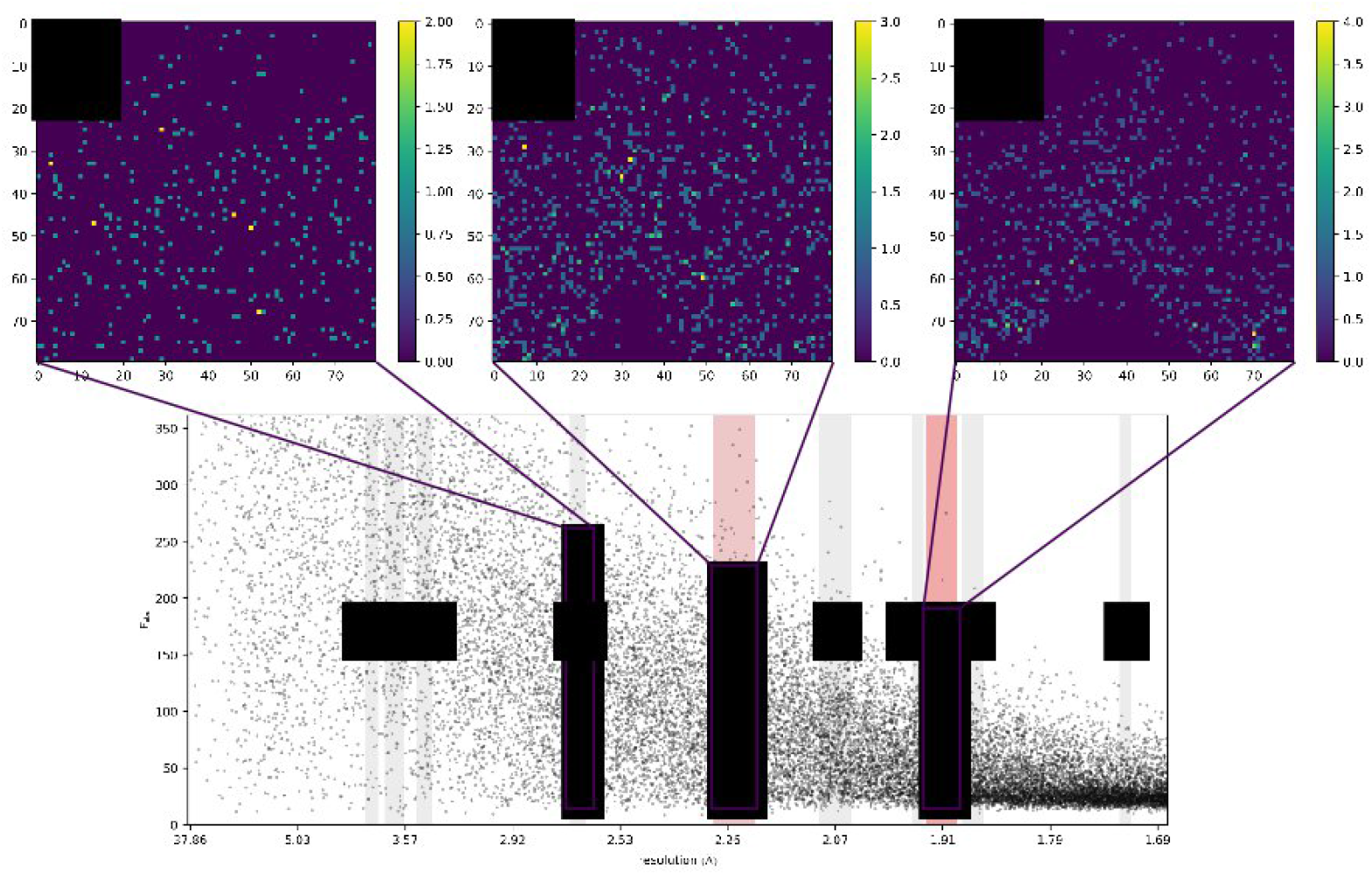
Relationship between the AUSPEX plot (bottom) (PDB-ID: 4epz) and the Helcaraxe plots (top). The grey bars in the AUSPEX plot indicate the resolution ranges in which an ice ring can appear and that are used to generate the focused plots (top). The rectangles indicate the area which the plots display. Plot E and H show ice ring characteristics. Ice ring features were automatically identified by AUSPEX (red bars) by the vertical shift of *I*_obs_ or *F*_obs_ which occurs visually in the shape of a spike.

Identifying whether a structure - or more exactly the integrated, scaled and merged diffraction data set - available in the worldwide Protein Data Bank (wwPDB) (Berman, 2000) is affected by ice ring contamination is even more difficult. Only a few entries have the corresponding raw data (i.e. images) available, as these are neither required for publication nor can be deposited directly to the wwPDB. However, if an integrated and merged data set is affected by ice diffraction, one can assume that subsequent model refinement will be affected. It has been demonstrated that removing ice rings from the data during integration improves R values by as much as 4.8% (Parkhurst *et al*., 2017). Thus, the correct identification of ice rings in data sets is an important step in assessing and ultimately, improving data quality.

In addition to statistical identification in CTRUNCATE (Winn et al., 2011) and phenix.xtriage (Adams *et al*., 2010), the AUSPEX icefinder score, recently improved by Moreau and colleagues (Moreau *et al*., 2021), is one of the most reliable statistical tools to detect ice crystal artefacts in integrated, merged and scaled diffraction data sets. While statistical identification can identify stronger ice diffraction in processed data automatically, less distinct ice rings can go unnoticed. For this reason, AUSPEX also produces plots of observed intensities (*I*_obs_) (or structure factor amplitudes *F*_obs_) against resolution, which permit easy visual identification of ice ring contamination (Fig. 2) (Thorn *et al*., 2017).

**Figure 2.**
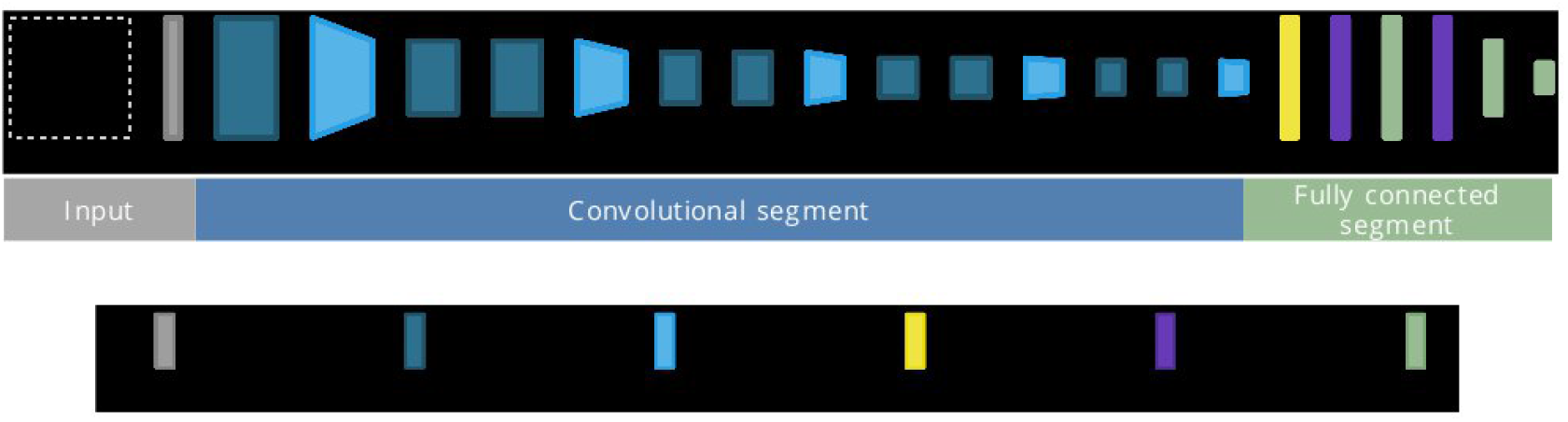
Schematic of the Helcaraxe network architecture. Input dimensions are vertically shown, the filter size horizontally. The blue part marks the convolutional segment while the fully connected part is green and purple. Image created with Net2Vis (Bäuerle & Ropinski, 2019).

The discrepancy that humans can easily recognize ice rings in these AUSPEX plots while automatic statistical detection remains difficult has led us to attempt an identification using artificial intelligence. In recent years, the use of Convolutional Neural Networks (CNNs) for data-driven research enabled the identification and recognition of complicated patterns in noisy data (Schmidt, 2019), leading to advances in all disciplines of science and data analysis. CNNs are exceptionally suited to classification of multi-dimensional arrays because they can retain spatial input information (Yamashita *et al*., 2018). Here, we present the results of employing CNNs to detect ice artefacts in processed macromolecular diffraction data.

## 2. Methods

### 2.1. Selection of training, validation and test data

1827 integrated, scaled, and merged diffraction data sets indicated to have been measured at 100 K were used to generate training and validation sets (Supporting Information: Helcaraxe_train_labels.xlsx). These diffraction data were randomly selected from the Coronavirus Structural Task Force repository (Croll et al., 2021) (396 diffraction data sets), the Integrated Resource for Reproducibility in Macromolecular Crystallography (Grabowski et al., 2016) (280) and the Protein Data Bank (1151) (Berman, 2000) without duplicates. Diffraction data were used in MTZ format, obtained through the conversion of sf.cif files by CCP4 cif2mtz (Winn et al., 2011). If CIF files had no observed Intensities (*I*_obs_), structure factor amplitudes (*F*_obs_) were used instead. If MAD data had been measured, the wavelength listed first in the deposited CIF file was used.

To convert the data into a format that could be presented to a neural network two-dimensional histograms of *I*_obs_ or *F*_obs_ against resolution were generated, dubbed “Helcaraxe plots”, using the NumPy *histogram2d* function (Virtanen *et al*., 2020) (example code in Supporting Information). The size of the histograms was set to 80×80 pixel as this has proven to be the best compromise between information loss and data size. Multiple Helcaraxe plots were produced per dataset, one around every expected ice-ring position present in the overall resolution range of the diffraction data set (Fig. 2 top). The width of individual ice ring resolution ranges has been previously identified (Thorn *et al*., 2017). To avoid extreme intensity outliers and for normalisation, the lower limit of the y-axis was set at the 0.5^th^ percentile of intensities or amplitudes and the top limit at the 95^th^ percentile. These parameters have proven to be the best middle ground between data loss and plot similarity. The x-axis was scaled to obtain constant histogram size despite the different widths of the individual ice ring resolution ranges.

80% of Helcaraxe plots generated from the MTZ files were allocated randomly to the training set and 20% to the validation set. The validation set was used to evaluate the CNNs during training and to select two final CNN candidates for the Helcaraxe program. The test set was not used in training:

A set of 200 randomly chosen diffraction data sets labelled for ice ring contamination (previously published as AUSPEX Appendix section C and reproduced here in Supplementary Information), was assigned as a test set (Thorn *et al*., 2017).

### 2.2. Data annotation

For annotation, ice rings were first identified in diffraction data sets by AUSPEX plots. Subsequently, Helcaraxe plots for training, validation, and test sets were generated as described and manually annotated for ice ring presence using the previous annotation results as guidance. A Helcaraxe plot was labelled as ice diffraction contaminated if at least two of the following criteria were met:

I. A vertical shift of *I*_obs_ or *F*_obs_ values in the shape of a spike must be visible to the naked eye.
II. At least 1% of the area of the plot was affected by the ice ring, meaning that either a part of the plot was blank because of the ice ring or intensities were shifted upwards. The area was measured by overlaying a grid.
III. The ice ring must be visible in the corresponding AUSPEX plot. 486 (26.6%) of the 1827 diffraction data sets used for the training and validation set were found to have ice diffraction contamination according to the aforementioned criteria (see Table 1). This resulted in the generation of 13,170 individual Helcaraxe plots, of which 984 (7.47%) were annotated as contaminated (Supplementary Information: Helcaraxe_train_labels.xlsx).

**Table 1.** Composition of training, validation, and test sets, divided into F_obs_ and I_obs_. The percentages of Helcaraxe plots showing ice ring contamination are given in brackets.

|  | Training | Validation | Test |
| --- | --- | --- | --- |
| **$F_{\text{obs}}$** | **6849** | **1713** | **1032** |
| *No ice contamination* | 6251 | 1564 | 977 |
| *Ice contamination* | 598 (8.73 %) | 149 (8.70 %) | 65 (6.24 %) |
| **$I_{\text{obs}}$** | **3686** | **922** | **1410** |
| *No ice contamination* | 3497 | 874 | 1329 |
| *Ice contamination* | 189 (5.13 %) | 48 (5.21 %) | 81 (5.74 %) |

The test set (AUSPEX Appendix section C), which was previously labelled for ice ring contamination, includes ice ring classification from other software and was also used by Moreau and colleagues to compare the performance of their algorithm (Moreau *et al*., 2021). The test set was again reviewed where Moreau’s annotation was consulted, and 9 labels (4.5%) were updated. (Supporting Information: Helcaraxe_test_results.xlsx). Three diffraction data sets were omitted because they were already superseded or had a lower maximum resolution than those ranges contaminated by ice rings. Of the 197 diffraction sets 40 (20.3%) were labelled as containing an ice ring. There was no overlap between the training/validation and the test set.

### 2.3. Network architecture and training

The network architecture of the employed CNNs consists of a convolutional and a fully connected part (Fig. 3). Two networks with the same architecture were trained, one for Iobs and one for Fobs values. Helcaraxe plots were supplied through an input layer that passes the plot directly to the subsequent convolutional segment. The first segment of the network extracts data features and consists of four blocks, with each block having two convolutional layers followed by an aggregating MaxPooling layer (Fig. 4) and Batch Normalization layers in between to reduce the risk of overfitting through normalization. The second segment is connected through a Flatten layer and contains two fully connected layers of artificial neurons separated by dropout layers which randomly omit neurons during training to further reduce the risk of overfitting (Srivastava et al., 2014). The output layer is a single neuron that uses a sigmoid activation function. Therefore, a value (hereafter referred to as prediction) between 0 (no ice diffraction artefacts) and 1 (ice diffraction artefacts) is returned for each Helcaraxe plot. The threshold for classification was 0.5.

**Figure 3.**
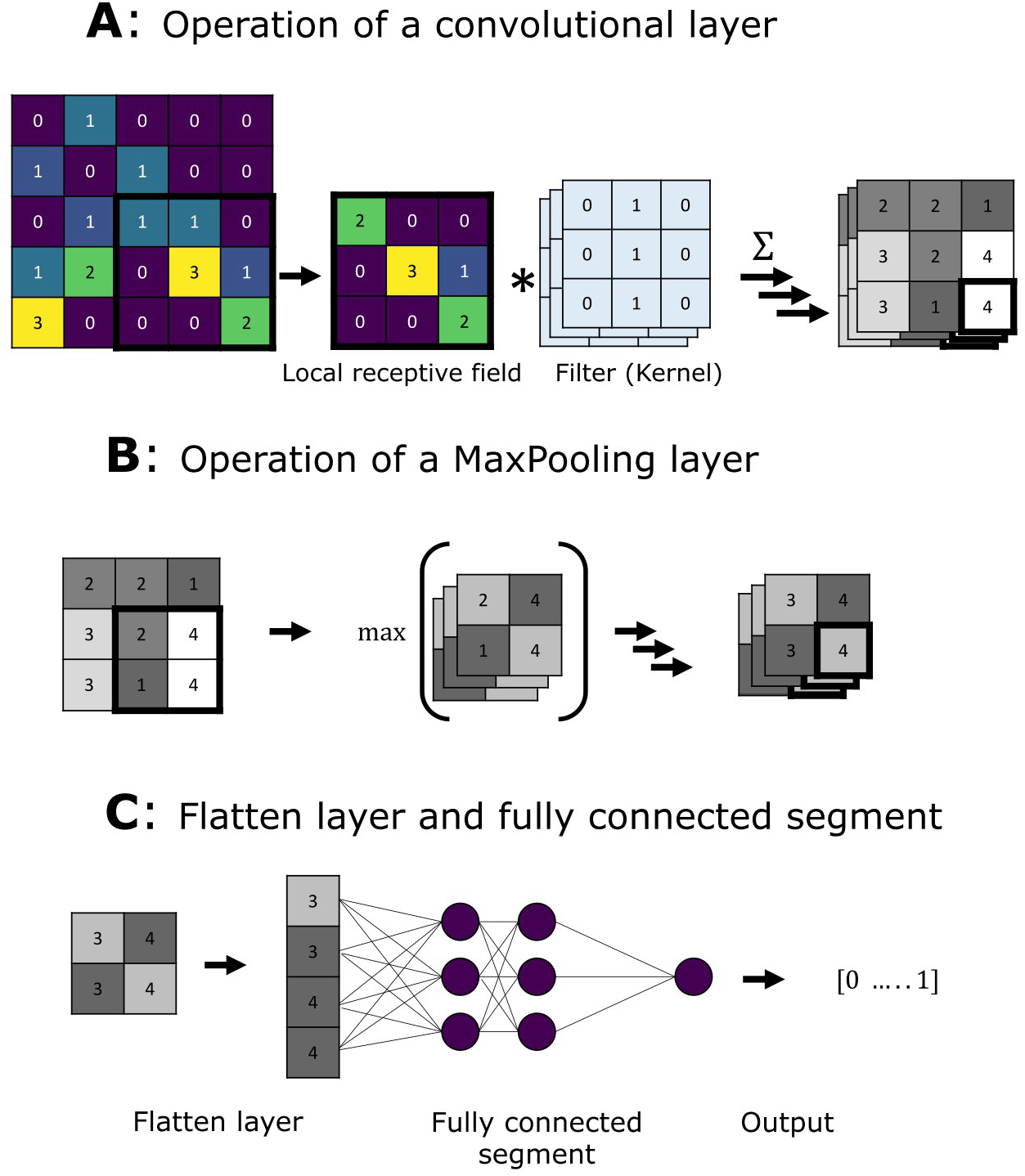
A: General operation of a convolutional layer to create a feature map. A patch (the local receptive field) of the input array is altered through multiple filters and the resulting values are saved in the feature map. B: Like a convolutional layer, the pooling layer is only connected to a limited group of inputs within a rectangular field. However, it has no filter and only summarizes the input matrix through aggregation. C: The flatten layer flattens the multidimensional matrix into a single-dimensional one and the output is passed to a regular neural network made of fully connected artificial neurons.

**Figure 4.**
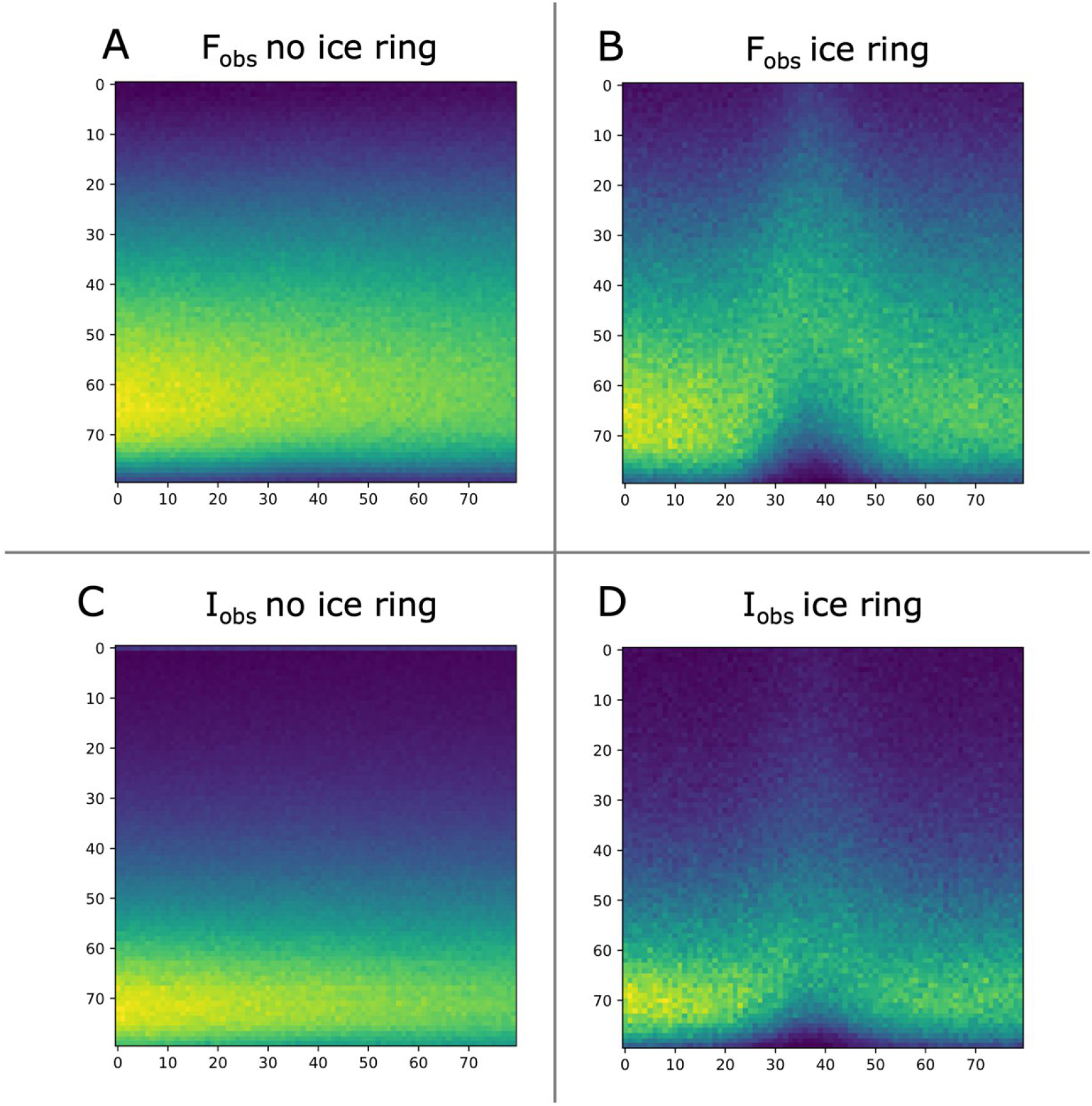
Averaged Helcaraxe plots of training and validation data, annotated as containing or not containing ice artefacts, for both, *F*_obs_ and *I*_obs_. The characteristic ice spike can be seen in both mean ice ring plots (B and D). In *F*_obs_ the spike is more prominent.

Parameters which control the learning process (hyperparameters) were optimized using the Hyperband optimization algorithm (Li *et al*., 2018). The network used for predicting ice diffraction *F*_obs_ plots (hereafter referred to as *F*_obs_ network) was trained and validated only using *F*_obs_ Helcaraxe plots, the network for predicting *I*_obs_ plots (hereafter referred to as *I*_obs_ network) was trained through transfer learning, fine-tuning the *F*_obs_ network using *I*_obs_ plots and a very moderate rate of learning (0.0005). This was done to make sure that the network could adapt to the differences in the *I*_obs_ Helcaraxe plots without overriding the pattern recognition abilities already acquired by the *F*_obs_ network.

Network design and training were performed using TensorFlow 2.4.1 (Abadi *et al*., 2016). The final trained networks were selected from multiple training runs based on the performance against the validation set. Their ability to operate reliably on unseen data was tested using the independent test set.

## 3. Results

### 3.1. Data features

To acquire an overview of how ice diffraction artefacts manifest in Helcaraxe plots, all plots from the training and validation sets which manually had been annotated as not containing ice diffraction were averaged, for amplitudes and intensities, respectively (Fig. 5A and C), as were all plots annotated as containing ice diffraction (Fig. 5B and D). The resulting averaged plots of intensities or structure factor amplitudes with no ice show a uniform vertical gradient. The corresponding plots with ice artefacts show an upward shift in the form of a spike in the middle. It is apparent that the points spread more evenly in *F*_obs_ than in *I*_obs_ plots. Spikes are also more visually prominent in Fig. 5 B than D, potentially a consequence of conversion of intensities into amplitudes by the French & Wilson method (French & Wilson, 1978), which imposes distribution expectations in order to facilitate conversion.

**Figure 5.**
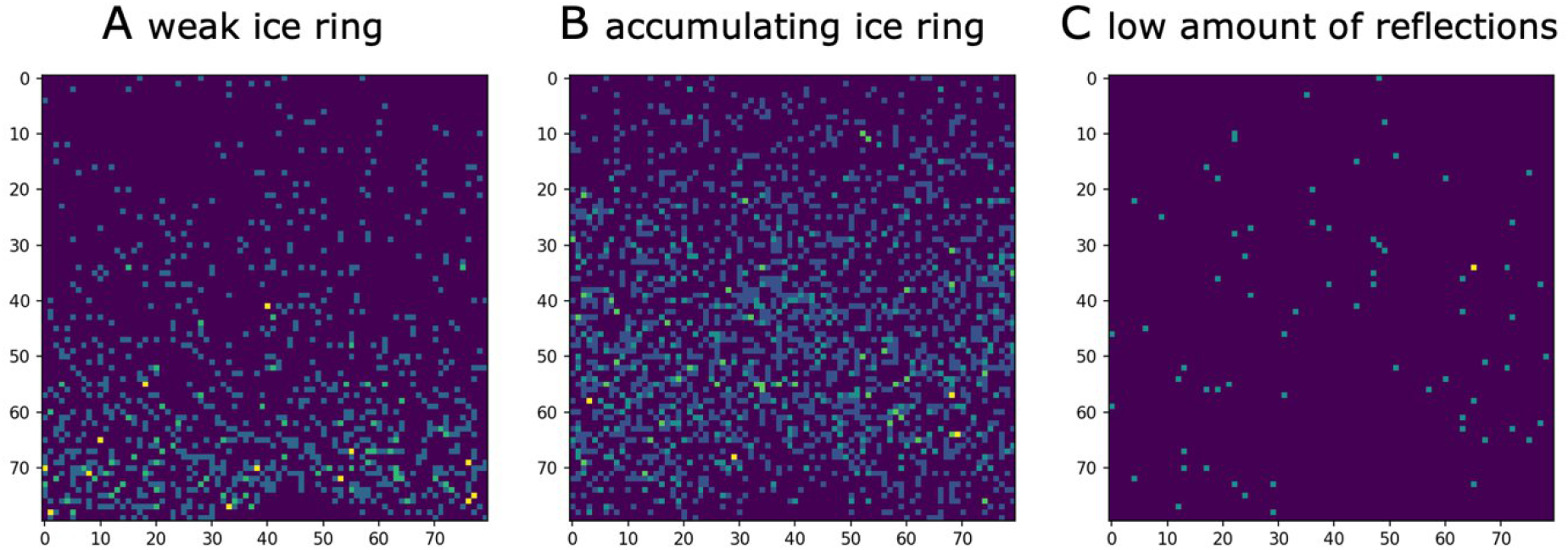
Misclassified plots. Weak (A) “accumulating” (B) ice rings in plots were the main causes of false negative classification. “Accumulating” refers to ice rings which do not affect all intensities equally at a given resolution. Diffraction data with a low number of reflections (C) was a source of false positive misclassification.

### 3.2. Network performance

Two trained networks were selected from multiple training runs based on their performance on the validation set. They were both evaluated against the test set (see section 3.2) to confirm that they can generalize. The performance was measured using three metrics: accuracy (1), sensitivity (2) and specificity (3):

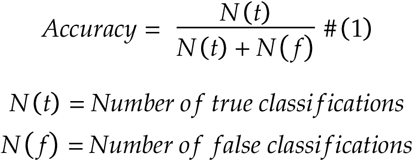

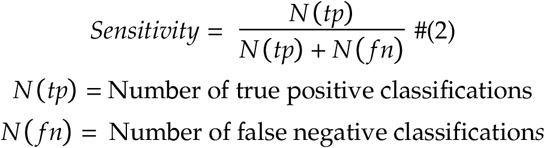

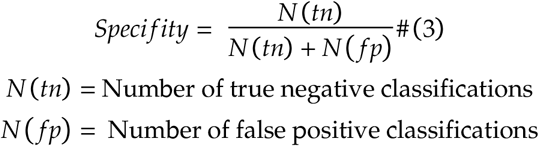

Judging by these criteria, the networks perform well on the validation set used in network training and the independent test set (see Table 2), showing there was no overfitting of the networks with regard to the training set. The performance of both networks is sufficient to detect ice-rings in most cases. Both networks have a higher specificity than sensitivity.

**Table 2.** Performance of the two final *F*_obs_ and *I*_obs_ networks against the Helcaraxe plots of the validation and test set. Accuracy, sensitivity and specificity values are given with a 95% confidence interval.

|  | True<br>positive | False<br>positive | True<br>negative | False<br>negative | Accuracy | Sensitivity | Specificity |
| --- | --- | --- | --- | --- | --- | --- | --- |
| $F_{\text{obs}}^{\text{validation}}$ | 92 | 8 | 1556 | 57 | 0.962 0.009 | 0.617 0.003 | 0.995 0.0001 |
| $F_{\text{obs}}^{\text{test}}$ | 42 | 4 | 973 | 13 | 0.984 0.007 | 0.764 0.004 | 0.996 0.0001 |
| $I_{\text{obs}}^{\text{validation}}$ | 37 | 6 | 868 | 11 | 0.982 0.009 | 0.771 0.003 | 0.993 0.0001 |
| $I_{\text{obs}}^{\text{test}}$ | 69 | 13 | 1316 | 12 | 0.982 0.007 | 0.852 0.002 | 0.990 0.0002 |

**Table 3.**
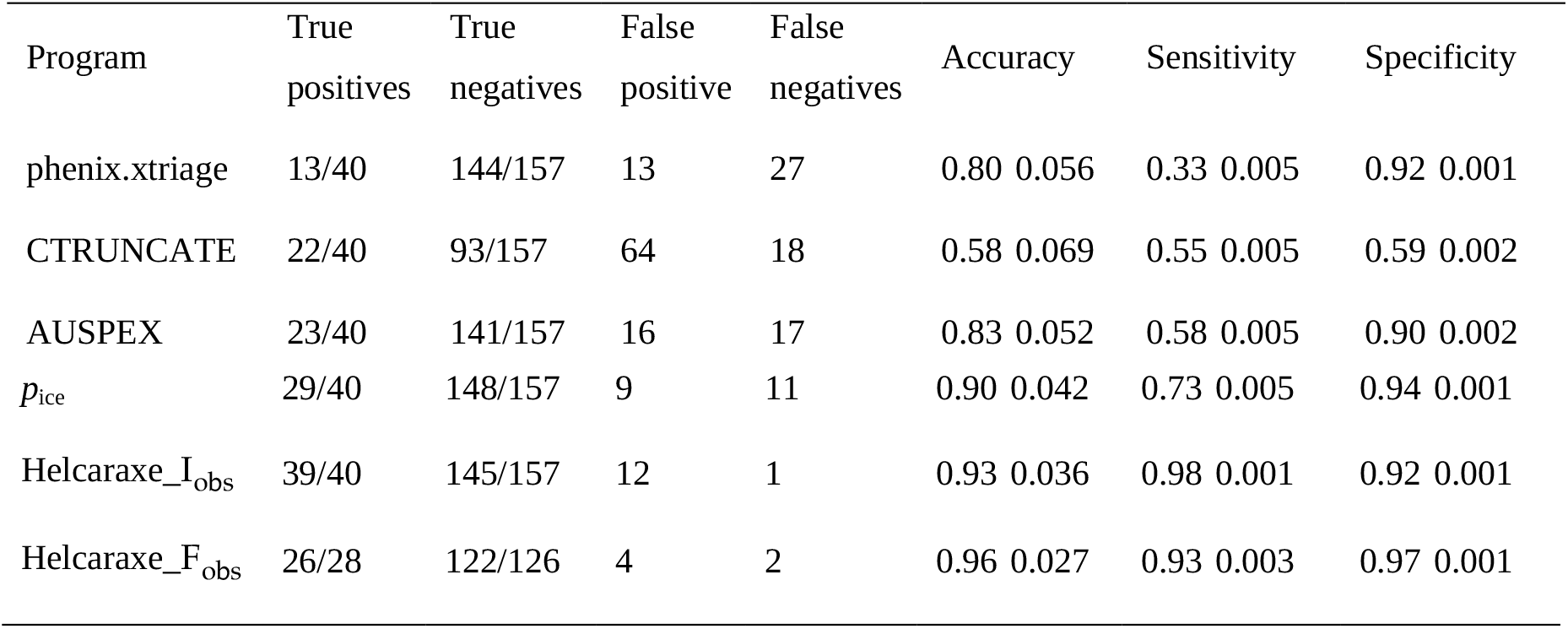
Comparison of different ice detecting software. *p*_ice_ is the new score introduced by Moreau and colleagues based on the previous AUSPEX icefinder score. The data of accuracy, sensitivity and specificity are given with a 95% confidence interval.

Closer visual inspection of false negative classifications reveals that a portion of all test plots in the test set, falsely identified as negative, had only very small spikes affecting approximately 1% of the plot (as described in definition II in 2.1) (*I*_obs_: 7 of the 12 false negative classifications) (Fig. 6A) or the diffraction data set contained relatively few reflections (Fig 6C) (*I*_obs_:5 of 12, *F*_obs_: 2 of 13). Another cause of false negative classification is the presence of data points in the area of the ice spike (*I*_obs_:4 of 12, *F*_obs_: 8 of 13) (Fig. 6C). We suspect this occurs when the ice crystals build up during measurement so that both contaminated and uncontaminated intensities are present in the merged diffraction data. The main reason for false positive classification was shift or absence of intensities in the usual ice ring range without the typical shape of a spike (as described in definition I in 2.1) (*I*_obs_: 7 of the 13 false positive classifications, *F*_obs_: 2 of 4).

**Figure 6.**
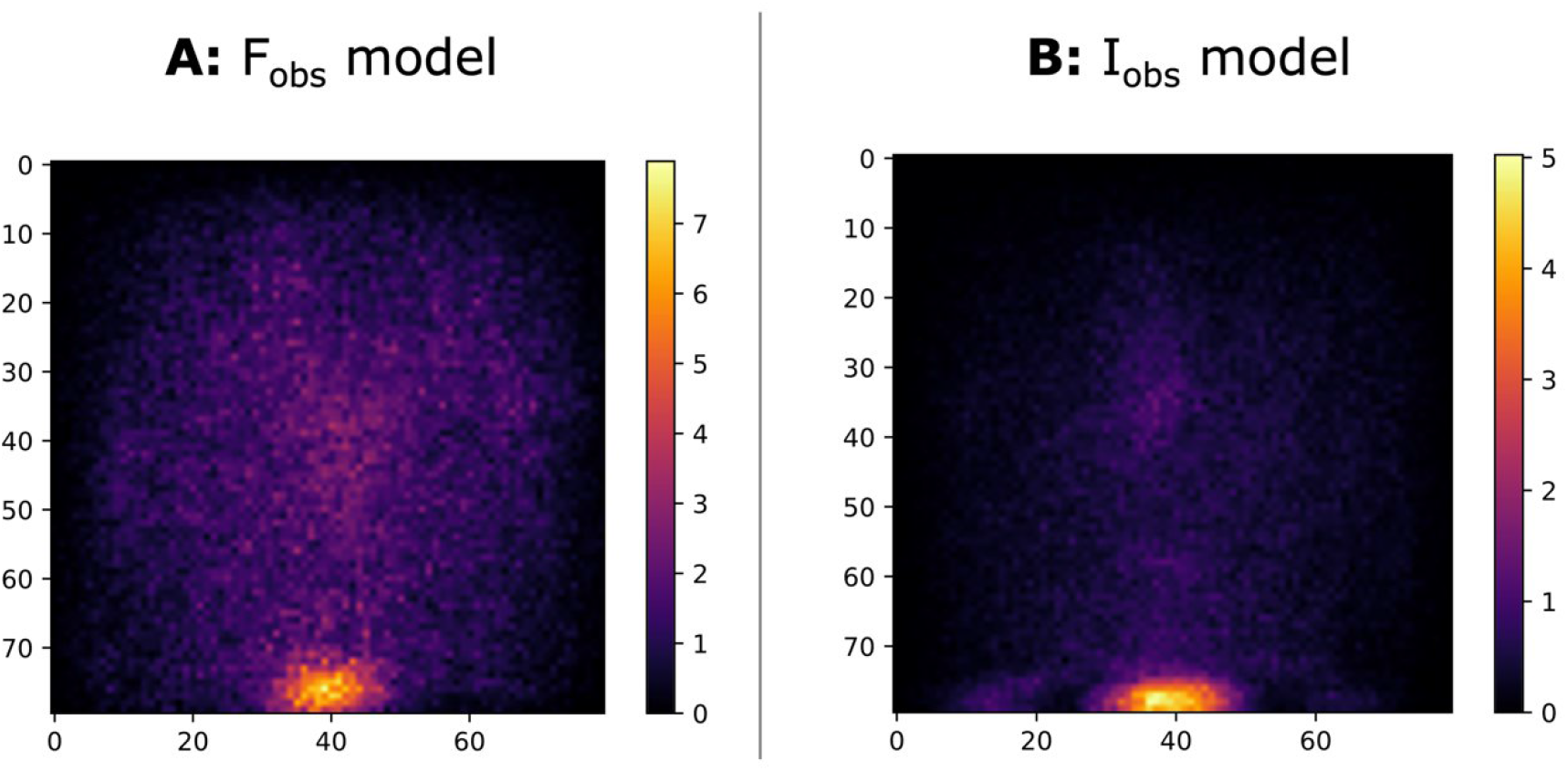
SmoothGrad’s averaged sensitivity mask of the *F*_obs_ and *I*_obs_ network against the test set. Larger values indicate a higher significance of the pixel. The area where ice artefacts usually appear have higher relevance than the top, left and right edges. *F*_obs_ plots have a more spread out distribution, which is the likely reason why the *F*_obs_ network has also a larger relevant area.

To obtain insight into the decision-making process of the networks, SmoothGrad (Smilkov *et al*., 2017) was used to generate sensitivity maps that highlight which area has the most impact on the classification of the Helcaraxe plot. The area at the bottom, especially in the middle (where ice rings usually appear) had the greatest influence on the decision of the network. The edges on the top, right, and left had close to no impact on the classification. Fig. 7 shows that the networks recognize the characteristic ice spike in Helcaraxe plots and uses it as indicator for the classification. The comparison of the two sensitivity maps suggests that the two models have adapted to the properties (as described in 3.1) of their respective Helcaraxe plots (as described in 3.1). *F*_obs_ plots have a more spread-out distribution than *I*_obs_ and the sensitivity map shows the Fobs model is sensitive to a broader area.

**Figure 7.**
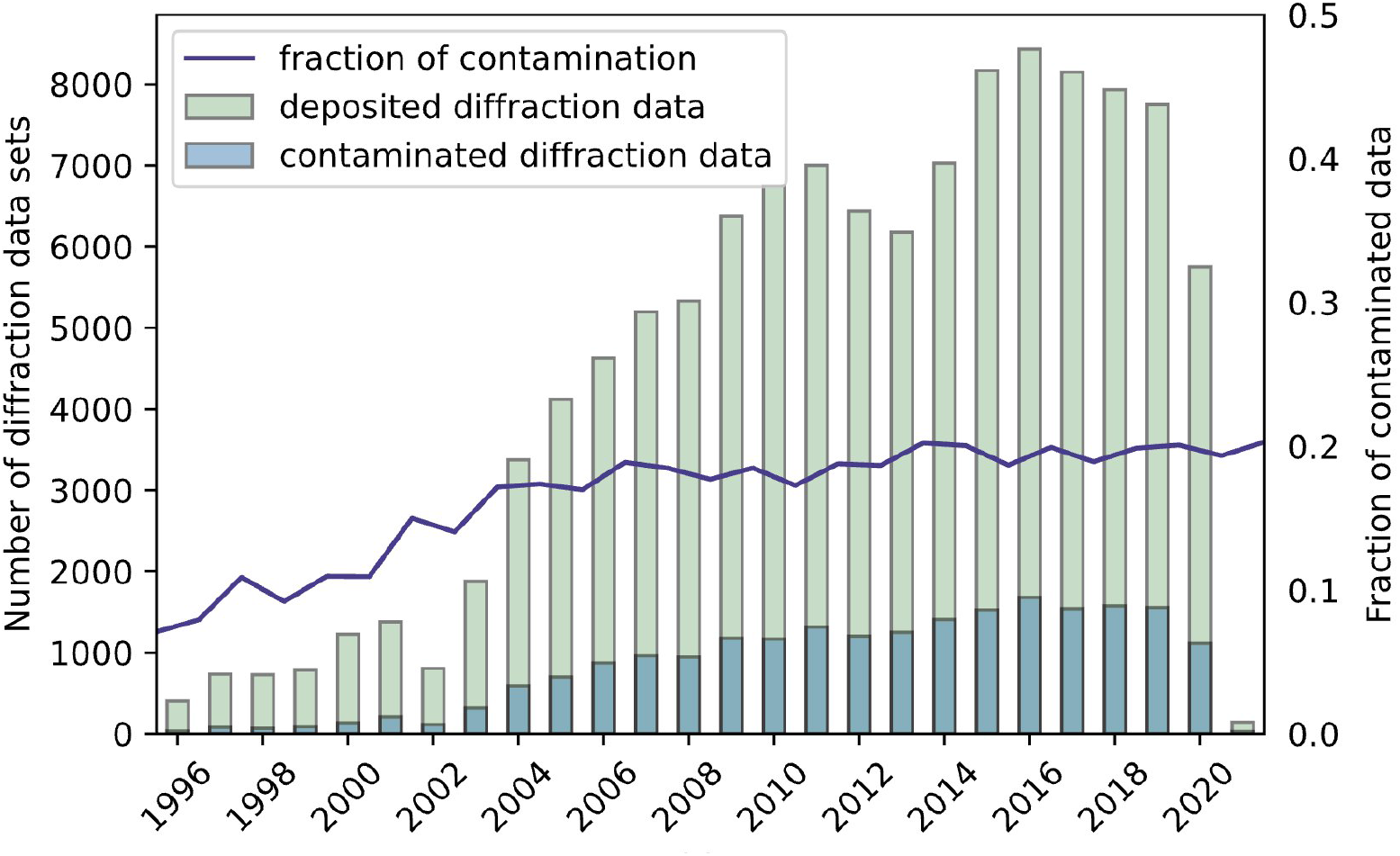
Number of deposited diffraction data sets (green) and the amount of these depositions which were annotated by Helcaraxe as containing at least one ice ring (blue). The contamination proportion is shown as a line (purple).

### 3.3. Comparison to other algorithms

Performance of both *F*_obs_ and *I*_obs_ networks against the test set was compared to other ice ring detection algorithms, namely phenix.xtriage (Adams *et al*., 2010), CTRUNCATE (Winn *et al*., 2011), the AUSPEX icefinder score and the recent *p*_ice_ algorithm (Moreau *et al*., 2021). Helcaraxe rates the individual resolution ranges in which ice rings can appear (as described in 3.1) and not the complete diffraction data set. Therefore, a data set was labelled as ice ring contaminated when the network classified even a single Helcaraxe plot as contaminated.

The algorithm recently introduced by Moreau and colleagues outperformed all previous statistical methods. Helcaraxe reaches an even higher accuracy and sensitivity (both over 93%). Using *I*_obs_ plots, Helcaraxe has a specificity of 92%, using *F*_obs_ plots 97%. In general, the use of Helcaraxe results in the most reliable output in comparison to the traditional statistical methods.

### 3.4. Analysis of the PDB

The Helcaraxe networks were then used to analyse 117,615 randomly selected diffraction data sets deposited in the PDB. We found that 21,741 (18.5%) PDB entries show evidence of ice contamination in the processed, scaled, and merged diffraction data. This number is similar to other previous large-scale analyses of the PDB (19%, Thorn et al., 2017, 16 % Moreau et al.). We analysed the historical evolution of the ice contamination. Ice ring presence started rising in the late 1990s when cryo techniques became routine at the first synchrotron MX beamlines and steadily grew when the first sample changer came online. Since the mid-2000s, the fraction of ice contaminated data has stabilized around 19%, even though the advent of pixel detectors translated into shorter measurement times and consequently, less time in which the protein crystal is exposed to a cryo-stream where it may accumulate surface ice crystals on the sample. The data produced in this analysis could be a useful starting point for the research into the impact ice rings have on structure solution and will be available from Helcaraxe’s git repository (https://github.com/thorn-lab/helcaraxe).

### 3.5. Integration into AUSPEX

Helcaraxe has been integrated into a new Python-based version of AUSPEX (to be published) and can be used to automatically decide whether a specific resolution range contains ice ring artefacts. The previous icefinder score (Thorn *et al*., 2017) is still used to rate the severity of the artefacts as Helcaraxe, trained only for the detection and not the assessment of ice rings, provides a less differentiated classification. An additional discriminator function in the Helcaraxe script detects plots that are completely or partially blank, for example, because resolution ranges were omitted during integration. This is achieved by checking whether, for specified resolution bins, the mean *I*_obs_ (or *F*_obs_) is close to 0. If this is true, the plot is marked as non-predictable, and a warning is passed to the user. The runtime of AUSPEX is barely influenced by Helcaraxe as the prediction is fast (1 −3 seconds per diffraction data set). No additional input for AUSPEX is needed to use Helcaraxe, and users of AUSPEX can choose between the Helcaraxe networks or AUSPEX icefinder score to detect the potential ice crystal artefacts.

## 4. Conclusion

The identification of ice diffraction artefacts in integrated, scaled and merged data has been an ongoing problem in macromolecular crystallography, even with modern cryo-cooling techniques (Moreau *et al*., 2021) and new background estimation algorithms (Parkhurst *et al*., 2017). To aid identification in automatic pipelines as well as by users, a set of neural networks named Helcaraxe was developed to identify whether a scaled and merged X-ray diffraction dataset contains ice diffraction contamination of the reflection data from a macromolecular crystal. This program presents a significant improvement over previous automatic tools using classical statistical indicators. One area of future exploration would be to combine these approaches: Reliable statistical methods such as those recently introduced by Moreau and colleagues could be used as an additional feature in the fully connected part of Helcaraxe’s networks. Our work also shows that the multi-dimensional pattern recognition abilities of convolutional neural networks are a valuable addition to the toolbox of diffraction data analysis and the authors of this paper expect to see a rise of AI methods in this field in the near future. Helcaraxe is currently already in use in the Coronavirus Structural Task Force pipeline (Croll et al., 2021) and is integrated into the newest AUSPEX version which is available through the AUSPEX webserver (https://www.auspex.de).

## Acknowledgements

This work was supported by the German Federal Ministry of Education and Research [grant no. 05K19WWA] and the Deutsche Forschungsgemeinschaft [grant no. TH2135/2-1] The authors would like to thank Robert Nicholls, Gianluca Santoni, Maximilian Edich, Ferdinand Kirsten, Luise Kandler, Matt Reeves and Arwen Pearson for discussion.

## Notes

### Competing Interest Statement

The authors have declared no competing interest.

## References

Abadi, M., Agarwal, A., Barham, P., Brevdo, E., Chen, Z., Citro, C., Corrado, G. S., Davis, A., Dean, J., Devin, M., Ghemawat, S., Goodfellow, I., Harp, A., Irving, G., Isard, M., Jia, Y., Jozefowicz, R., Kaiser, L., Kudlur, M., Levenberg, J., Mane, D., Monga, R., Moore, S., Murray, D., Olah, C., Schuster, M., Shlens, J., Steiner, B., Sutskever, I., Talwar, K., Tucker, P., Vanhoucke, V., Vasudevan, V., Viegas, F., Vinyals, O., Warden, P., Wattenberg, M., Wicke, M., Yu, Y. & Zheng, X. (2016). ArXiv:1603.04467 [Cs].

Adams, P. D., Afonine, P. V., Bunkóczi, G., Chen, V. B., Davis, I. W., Echols, N., Headd, J. J., Hung, L.-W., Kapral, G. J., Grosse-Kunstleve, R. W., McCoy, A. J., Moriarty, N. W., Oeffner, R., Read, R. J., Richardson, D. C., Richardson, J. S., Terwilliger, T. C. & Zwart, P. H. (2010). Acta Crystallogr D Biol Crystallogr. 66, 213–221.

Bäuerle, Alex, Christian van Onzenoodt, and Timo Ropinski.” arXiv e-prints (2019): arXiv-1902.

Berman, H. M. (2000). Nucleic Acids Research. 28, 235–242.

Chapman, M. S. & Somasundaram, T. (2010). Acta Crystallogr D Biol Crystallogr. 66, 741–744.

Croll, T., Diederichs, K., Fischer, F., Fyfe, C., Gao, Y., Horrell, S., Joseph, P., Kandler, L., Kippes, O., Kirsten, F., Müller, K., Nolte, K., Payne, A., Reeves, M. G., Richardson, J., Santoni, G., Stäb, S., Tronrud, D., Williams, C. & Thorn, A. (2021). Nature Structural & Molecular Biology. 11.

Czyzewski, A., Krawiec, F., Brzezinski, D., Porebski, P. J. & Minor, W. (2021). Expert Systems with Applications. 174, 114740.

French, S. & Wilson, K. (1978). Acta Cryst A. 34, 517–525.

Garman, E. F. & Owen, R. L. (2006). Acta Crystallographica Section D: Biological Crystallography. 62, 32–47.

Garman, E. F. & Weik, M. (2019). J Synchrotron Rad. 26, 907–911.

Grabowski, M., Langner, K. M., Cymborowski, M., Porebski, P. J., Sroka, P., Zheng, H., Cooper, D. R., Zimmerman, M. D., Elsliger, M.-A., Burley, S. K. & Minor, W. (2016). Acta Crystallogr D Struct Biol. 72, 1181–1193.

Li, L., Jamieson, K., DeSalvo, G., Rostamizadeh, A. & Talwalkar, A. (2018). Journal of Machine Learning Research. 18, 1–52.

Moreau, D. W., Atakisi, H. & Thorne, R. E. (2021). Acta Crystallogr D Struct Biol. 77, 540–554.

Parkhurst, J. M., Thorn, A., Vollmar, M., Winter, G., Waterman, D. G., Fuentes-Montero, L., Gildea, R. J., Murshudov, G. N. & Evans, G. (2017). IUCrJ. 4, 626–638.

Schmidt, J., Marques, M. R., Botti, S., & Marques, M. A. (2019). npj Computational Materials, 5(1), 1–36.

Smilkov, D., Thorat, N., Kim, B., Viégas, F., & Wattenberg, M. (2017). arXiv preprint ArXiv:1706.03825.

Srivastava, N., Hinton, G., Krizhevsky, A., Sutskever, I., & Salakhutdinov, R. (2014). The journal of machine learning research, 15(1), 1929–1958.

Thorn, A., Parkhurst, J., Emsley, P., Nicholls, R. A., Vollmar, M., Evans, G. & Murshudov, G. N. (2017). Acta Crystallographica Section D: Structural Biology. 73, 729–737.

Virtanen, P., Gommers, R., Oliphant, T. E., Haberland, M., Reddy, T., Cournapeau, D., Burovski, E., Peterson, P., Weckesser, W., Bright, J., van der Walt, S. J., Brett, M., Wilson, J., Millman, K. J., Mayorov, N., Nelson, A. R. J., Jones, E., Kern, R., Larson, E., Carey, C. J., Polat, İ., Feng, Y., Moore, E. W., VanderPlas, J., Laxalde, D., Perktold, J., Cimrman, R., Henriksen, I., Quintero, E. A., Harris, C. R., Archibald, A. M., Ribeiro, A. H., Pedregosa, F. & van Mulbregt, P. (2020). Nat Methods. 17, 261–272.

Winn, M. D., Ballard, C. C., Cowtan, K. D., Dodson, E. J., Emsley, P., Evans, P. R., Keegan, R. M., Krissinel, E. B., Leslie, A. G. W., McCoy, A., McNicholas, S. J., Murshudov, G. N., Pannu, N. S., Potterton, E. A., Powell, H. R., Read, R. J., Vagin, A. & Wilson, K. S. (2011). Acta Crystallogr D Biol Crystallogr. 67, 235–242.

Yamashita, R., Nishio, M., Do, R. K. G. & Togashi, K. (2018). Insights Imaging. 9, 611–629.

